# Spontaneous neural activity in the developing ant olfactory system

**DOI:** 10.64898/2026.10.25.761666

**Authors:** Dominic D. Frank, Daniel J. C. Kronauer

## Abstract

Spontaneous neural activity is a pillar of mammalian neurodevelopment. In mammals, developing olfactory sensory neurons (OSNs) show spontaneous activity patterns that depend on the olfactory receptor (OR) they express. This activity is necessary for proper sorting of OSN axons into olfactory bulb glomeruli. This contrasts with the simpler olfactory system of the fruit fly *Drosophila melanogaster*, where antennal lobe glomeruli form without functional ORs. Compared to *Drosophila*, ants have a vastly expanded olfactory system and, unlike *Drosophila*, require functional ORs for proper development. This raises the possibility that ants, like mammals, rely on spontaneous neural activity. To test whether developing ant OSNs are spontaneously active, we performed live calcium imaging in translucent pupae of the clonal raider ant *Ooceraea biroi*, using a transgenic line expressing the genetically encoded calcium indicator GCaMP6s in OSNs. Developing *O. biroi* OSNs exhibit spontaneous calcium transients from the earliest stages of glomerulus formation, and features of this activity change over the course of metamorphosis. During early stages, activity is faster, varies widely between cells, and is uncorrelated across the OSN population. By mid-metamorphosis, activity has slowed, with longer calcium events and fewer peaks per cell. As pupae approach eclosion, a new mode appears, with rhythmic, sawtooth-shaped calcium events that synchronously engage OSNs in both antennae. This stage-dependent spontaneous activity suggests that the expanded ant olfactory system may use a developmental logic more akin to mammals than to *Drosophila*.

## Introduction

The development of nervous systems and neural circuits is a complex problem. In the olfactory system, for example, olfactory sensory neurons (OSNs), each of which expresses one of many olfactory receptors (ORs), have to find their correct projection neuron (PN) partners (1, 2). Connections between OSNs and PNs form in the primary olfactory center of the brain, the olfactory bulb in mammals and the antennal lobe (AL) in insects (3). Here, OSNs expressing the same OR converge onto the same globular structure, called a glomerulus, where they synapse with PNs (4, 5). This organization reflects a spatial map that represents odors in both mammals and insects (6–8). The similarities do not extend to the ORs themselves, however, which are evolutionarily unrelated. Mammalian ORs are G protein-coupled receptors (9, 10), whereas insect ORs are heteromeric ligand-gated cation channels composed of a variable tuning subunit and the obligate co-receptor Orco (11–13). How OSNs sort themselves and find their correct glomerulus has been studied primarily in mice (*Mus musculus*) and fruit flies (*Drosophila melanogaster*), yielding contrasting answers as to whether ORs contribute to this process.

In mice, ORs play key roles in matching OSNs to their appropriate targets in the olfactory bulb. Genetically changing the OR expressed by a given OSN redirects the cell’s axon to a different position in the bulb, indicating that OR identity contributes to glomerular position (14). One mechanism linking receptor identity to target choice involves spontaneous neuronal activity (15, 16). Different ORs give rise to distinct characteristics of intrinsic electrical activity during early development, and these activity signatures affect the expression of axon sorting molecules important for normal glomerulus formation (17). Consistent with this model, constitutive silencing or depolarization of young OSNs results in adult mice with disrupted glomerular maps (17, 18).

In contrast, the fly AL develops normally under similar experimental perturbations (19). In flies, OSN identity is tied to a transcription factor cascade that regulates the expression of cell-surface proteins (20, 21). Together, these proteins constitute a combinatorial code for attraction and repulsion across cell types (22–25). During metamorphosis, this code guides OSN axons to their glomerular targets in the AL and matches them with the appropriate post-synaptic partners (26). Importantly, developmental progression is removed from the expression of molecular ORs in flies, and mutants lacking functional receptors still emerge from metamorphosis with ALs that are indistinguishable from wild type individuals (19, 27).

The olfactory system of *Drosophila* is much simpler than that of mammals, with ca. 62 ORs and 51 olfactory glomeruli per hemisphere compared to roughly 1,100 ORs and 2,800 olfactory bulb glomeruli in mice (28–30). This difference in complexity could help explain why the *Drosophila* olfactory system can be wired using a cascade of transcription factors, while the mouse relies on information from the actual ORs. However, the contrast between mammals and insects becomes less when considering a broader taxonomic scope. Ants, for example, usually possess several hundred ORs, and up to 687 ORs in *Pseudoneoponera rufipes* (31).These expansions are believed to be at least in part related to the evolution of the complex pheromone language that ants use to organize their societies (32–34). The OR numbers observed in ants overlap with mammalian OR repertoires, like the ∼400 functional ORs encoded in the human genome (35, 36), and they are reflected in similarly striking increases in the number of AL glomeruli (37–41).

Based on the assumption that the complexity of an olfactory system is reflected in its development, it has been hypothesized that ants may employ mechanisms more akin to mammals than to *Drosophila* (42). In ants, tuning ORs and Orco are expressed early in metamorphosis, before glomeruli begin to form, and the Orco protein is trafficked to OSN axon terminals as they invade the antennal lobe (27). These findings parallel olfactory bulb development in mice (43–45) and are consistent with the idea that ORs might play a role in axon targeting, either directly or indirectly. Most fly OSNs, on the other hand, switch on OR expression later in development, after antennal lobe formation is well underway, and tuning ORs and Orco are only found in OSN somata and dendrites, but not in axon terminals (46–48).

Despite these tantalizing initial findings, it remains unknown whether ant OSNs exhibit the spontaneous activity seen in developing mouse OSNs during metamorphosis, which could guide OSN targeting and AL formation. To address this question, we developed a non-invasive preparation for *in vivo* two-photon imaging of clonal raider ant pupae throughout metamorphosis. Using transgenic ants expressing the genetically encoded calcium indicator GCaMP6s in OSNs, this preparation allows us to acquire volumetric data from both antennae at subcellular resolution during hours-long imaging sessions.

## Results

### Imaging OSN activity in ant pupae

If spontaneous OSN activity contributes to the assembly of olfactory circuits, it should be detectable during olfactory system formation. In the clonal raider ant *Ooceraea biroi*, AL glomeruli form during a period of roughly 8 days, ranging from pupal day 2 (‘P2’) to pupal day 10 (‘P10’) of metamorphosis (27). In very early pupae, OSNs are immature, expressing Orco and a few ORs while not having formed olfactory glomeruli yet. By P2, most ORs are expressed, OSN axons have reached the nascent AL, and the first handful of glomeruli have formed. Glomerular number then increases over the following week, reaching the full adult complement of ∼500 glomeruli at P10. OR expression reaches adult levels at P14, at which point pupae have melanized (27). Adults eclose a few days later (**Fig. 1 A**).

**Figure 1.**
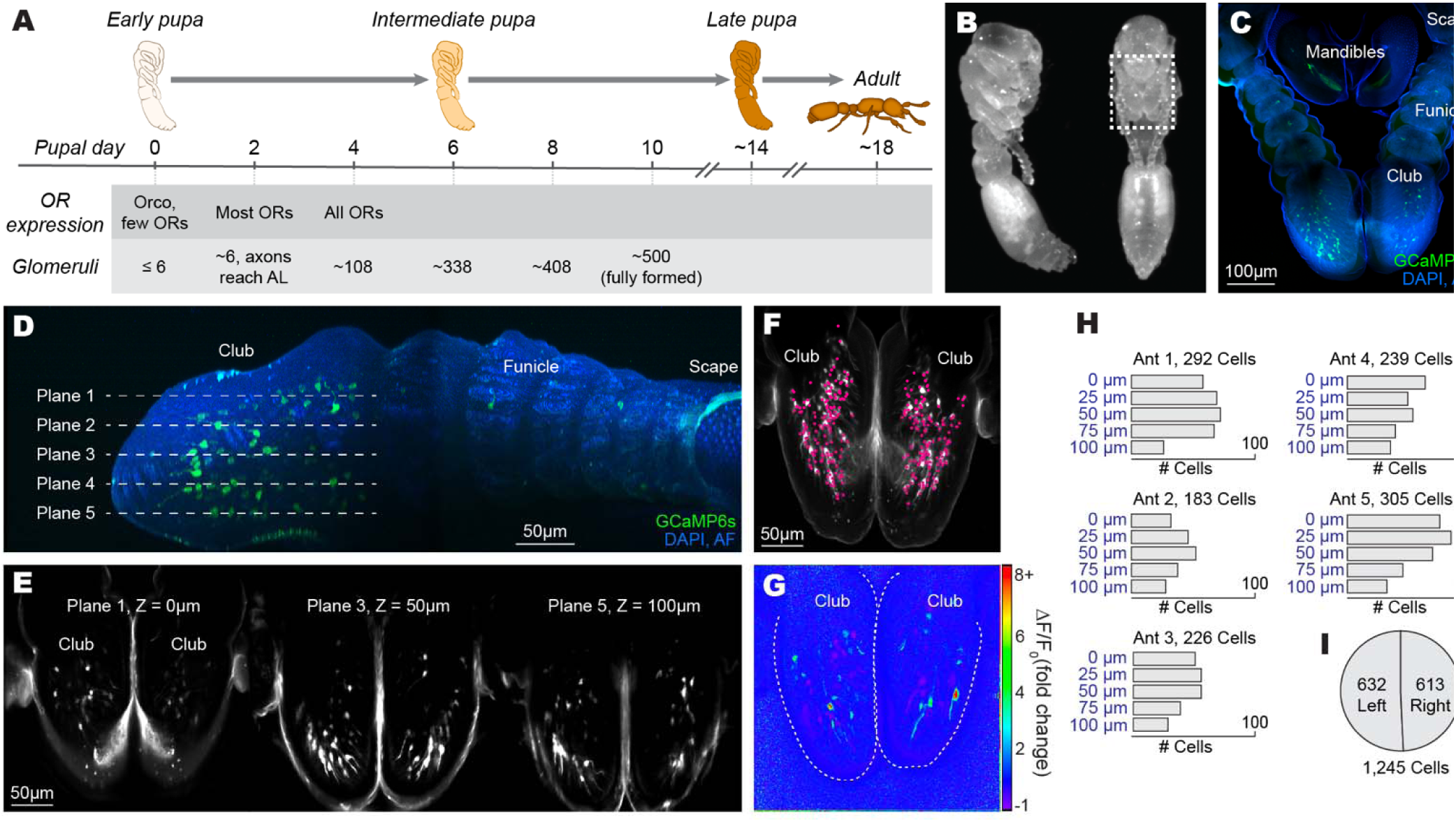
Imaging OSN activity in ant pupae. (A) Timeline of *Ooceraea biroi* olfactory system development, based on (27). (B) Photographs of an *O. biroi* pupa viewed from the side (left) and top (right). The white box indicates the location of the antennae. (C) Confocal micrograph of an early *O. biroi* pupa expressing GCaMP6s in antennal olfactory sensory neurons (OSNs, green). Nuclei are labeled with DAPI, and cuticular autofluorescence (AF) is visible in the same channel (blue). The field of view corresponds to the white box in (B). (D) Side view of the same antennae as in (C), with the approximate locations of two-photon imaging planes indicated. (E) Montage of representative frames from three example two-photon imaging planes spaced 50µm apart, starting from the most dorsal plane (far left) to the most ventral plane (far right). (F) Z-projection across all 5 imaging planes with the location of each manually segmented soma indicated by a magenta circle. (G) Representative ΔF/F image demonstrating spontaneous calcium transients in OSN somas and dendrites. The approximate perimeter of the antennal club is indicated by the dashed line. (H) Quantification of the number of OSNs recorded from each plane across five early pupae. (I) Pie chart quantifying the total number of OSNs recorded from the left and right antennae across these five individuals.

*In vivo* imaging of neural activity during insect metamorphosis has been challenging because in many species, including many ants, pupae are enclosed in a protective cocoon or puparium that hinders optical access. In the *D. melanogaster* pupa, for example, the hardened cuticle of the final larval instar forms a rigid and optically dense shell. While micro-dissections have allowed optical access to the brain via small windows (49), the peripheral chemosensory system in the antennae is particularly delicate, prohibiting the acquisition of activity data from developing chemosensory neurons. *O. biroi* pupae, however, are “naked”; they lack a cocoon or puparium, and their translucent cuticle provides direct optical access to internal structures (**Fig. 1 B**).Using an *O. biroi* transgenic line expressing GCaMP6s in OSNs under an *Orco* promoter fragment (*Orco>GCaMP6s;* **Fig. 1 C and D**) (50, 51), we exploited this feature to develop an *in vivo* volumetric two-photon imaging preparation where neural activity can be imaged simultaneously in both antennae at subcellular resolution and for extended periods of time **(Fig. 1 E)**.

We began our imaging experiments with P2 and P3 pupae (‘early’ pupae). At this age, our *Orco>GCaMP6s* line expresses in fewer cells than the endogenous *Orco* gene in wild-type ants (27). Antibody staining for Orco protein in transgenic ants confirmed that P2 OSNs express Orco broadly, as expected, suggesting that Orco expression in the transgenic line is normal and that the sparse GCaMP6s label is a technical artifact specific to this transgene (**Fig. S1 A**). Sparse labeling is advantageous in this case because it facilitates image segmentation. Using our *in vivo* preparation, we recorded GCaMP6s activity from all labelled OSNs at a rate of 0.8 seconds per volume and with each trial lasting 2 hours (**Fig. 1 E**). We then manually segmented OSN somata across all planes (**Fig. 1 F**) and calculated a ΔF/F timeseries for each cell (**Fig. 1 G**). These experiments resulted in a dataset of 1,245 neurons (249 +/- 50 neurons per pupa, n = 5 animals; **Fig. 1 H**), distributed roughly equally across the left and right antennae (**Fig. 1 I**). At this developmental stage, the *O. biroi* pupa has no visible olfactory sensilla and is enveloped in a fluid-filled membrane that prohibits exposure to the external environment (52). It is therefore unlikely that OSNs would come into contact with odorants in the air during imaging sessions.

### OSNs in early pupae show heterogenous and uncorrelated spontaneous activity

We observed a wealth of spontaneous calcium transients across the population, with a ΔF/F event of at least 1 (*i.e.,* a 100% increase) observed in 99.12% of recorded neurons (**Fig. 2 A**). Cells that did not reach this threshold were excluded from further analyses. Among the active cells, the features of the GCaMP traces varied dramatically: some neurons exhibited short, bursting calcium transients of smaller amplitudes, while others displayed minutes-long and large-amplitude calcium troughs and plateaus (**Fig. 2 B and C**). Estimating the number of ΔF/F peaks per cell and plotting this value against maximum ΔF/F revealed a modest linear relationship, where cells with many calcium peaks generally displayed smaller maximum peak amplitudes and those with few peaks displayed larger maximum amplitudes (r = −0.43, p < 0.001; R^2^ = 0.18) (**Fig. 2 D**).

**Figure 2.**
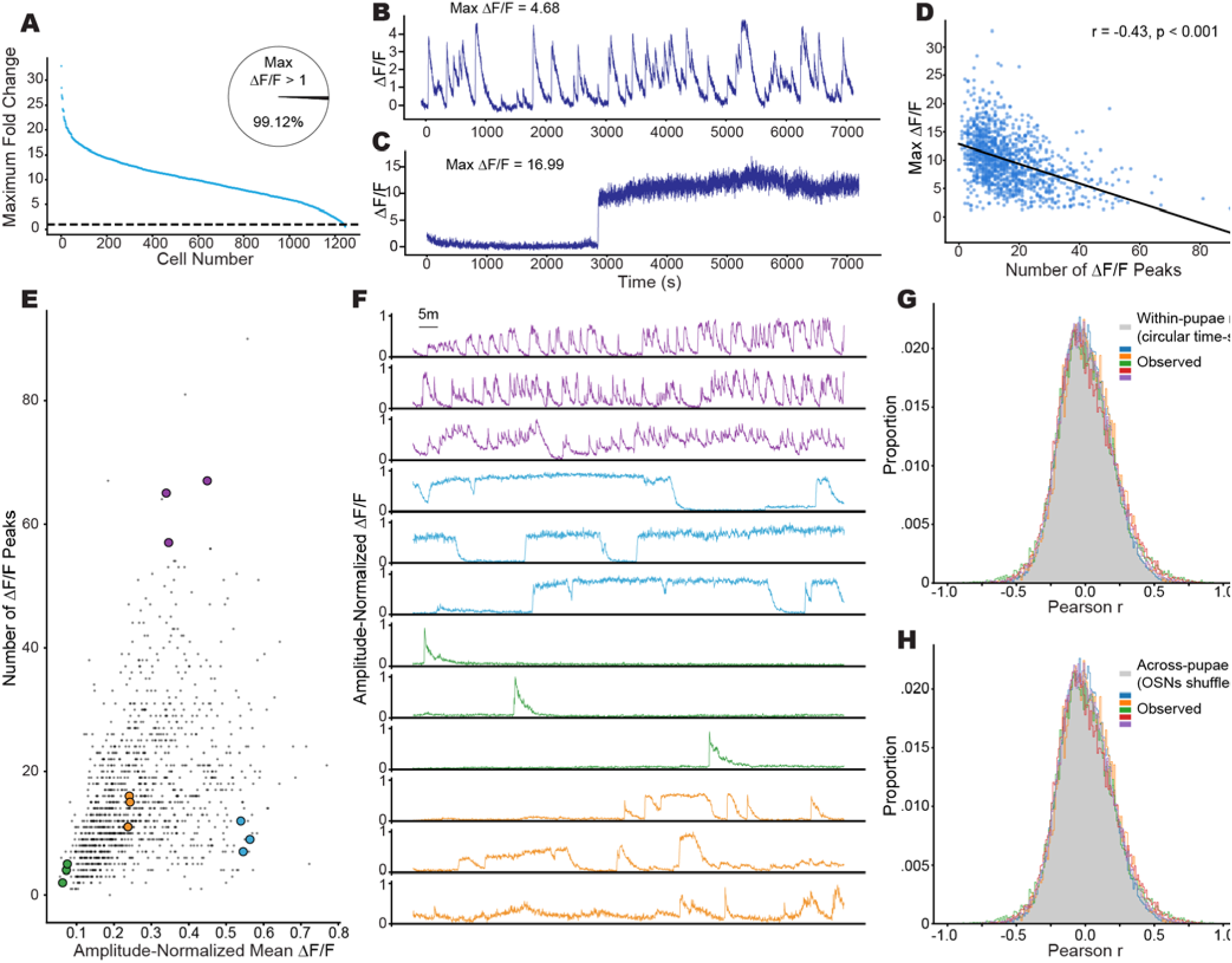
OSNs in early pupae show heterogenous and uncorrelated spontaneous activity. (A) Maximum ΔF/F fold change attained by each neuron in our dataset of 1,245 OSNs. Each point corresponds to a single neuron, and cells are sorted from largest to smallest maximum ΔF/F. The inset shows the percentage of neurons that exhibited at least one GCaMP transient exceeding 1 ΔF/F (*i.e.,* a 100% increase over baseline). (B-C) Two representative raw ΔF/F traces illustrating different temporal GCaMP activity types. (B) Shows a cell with frequent calcium transients and (C) shows a cell that stepped from the baseline to a stable plateau. (D) Linear regression plot of maximum ΔF/F and the number of ΔF/F peaks per active cell (n = 1,234 cells from n = 5 pupae). (E-F) Analyses of amplitude-normalized GCaMP activity. (E) Scatterplot comparing the number of ΔF/F peaks with the amplitude-normalized mean ΔF/F. Each data point is one neuron. (F) Representative time-series traces illustrating a diversity of different ΔF/F activity types. The colors of the traces correspond to the colored dots in (E). Traces are normalized and smoothed. (G-H) Pairwise correlations between simultaneously recorded OSNs. Histograms compare the observed distribution of Pearson correlation coefficients between OSN pairs to a null distribution generated by circularly shifting the traces in time (G) or shuffling the cells across pupae (H). Each colored line corresponds to the observed data from a different pupa.

Since GCaMP signal intensity could be influenced by expression differences across cells, we explored the calcium dynamics in our dataset independent of transient amplitude. We calculated an amplitude-normalized ΔF/F trace for every cell in our dataset and examined the relationship between the number of peaks and the temporal structure of the activity by plotting amplitude-normalized mean ΔF/F against the number of calcium peaks (**Fig. 2 E**). This revealed a continuous, wedge-shaped distribution with cells in different positions exhibiting different calcium dynamics (**Fig. 2 F**). For example, OSNs with many peaks and an intermediate mean ΔF/F exhibited rapid calcium fluctuations (**Fig. 2 F**, purple traces); cells with few peaks and large normalized mean ΔF/F displayed the long-duration calcium plateaus mentioned above (**Fig. 2 F**, blue traces); cells with few peaks and small mean ΔF/F displayed rare, brief calcium events (**Fig. 2 F**, green traces); and neurons with both an intermediate number of peaks and mean ΔF/F exhibited a mix of activity types (**Fig. 2 F**, orange traces). To test if cells recorded during the same session are temporally correlated with one another, we calculated a pairwise Pearson r for all cells from each animal and compared the distribution of values to a null distribution generated by circularly shifting each trace in time (within-pupae null, **Fig. 2 G**) or a null distribution created by pooling random cells from different animals (across-pupae null, **Fig. 2 H**). In both cases, the observed r values closely matched the null distributions, suggesting that calcium activity in early pupae is cell-autonomous and not strongly influenced by network effects or other extrinsic factors (*e.g.,* paracrine signaling). This is also consistent with the fact that OSN activity at this stage is spontaneous and intrinsic, rather than in response to external stimuli.

### OSNs in early pupae contain functional odorant receptors

In mammals, spontaneous GCaMP activity in developing olfactory sensory neurons is linked to the molecular olfactory receptor expressed by a given cell (*i.e.,* cells expressing different ORs exhibit different activity patterns) (17). Previous RNA sequencing studies have shown that OSNs in clonal raider ant pupae already express Orco and ORs at the developmental timepoint of our recordings (27), suggesting that molecular OR identity could also define activity patterns in our dataset. However, post-transcriptional regulation of ORs is abundant in the ant olfactory system (53–55), and the detection of OR transcripts alone does not demonstrate that functional Orco/OR channels are present in the membrane. We therefore asked whether Orco is properly localized to the membrane of OSNs in early pupae using immunohistochemistry and confocal microscopy. Compared to the cytosolic GCaMP6s signal, anti-Orco signal was observed in clear halos demarcating the boundaries of all olfactory sensory neurons (**Fig. S1 B and C**). This indicates that Orco/OR ion channels are indeed trafficked to and incorporated into the plasma membrane. Next, to confirm that these channels are indeed functional, we dissected antennae from early *Orco>GCaMP6s* pupae, bathed them in artificial hemolymph, and exposed them to the Orco agonist VUAA1. Introducing VUAA1 produced sustained, high-amplitude calcium transients across the OSN population, indicating that OSNs of P2 and P3 pupae indeed have functional odorant receptor ion channels (**Fig. S1 D**). This, together with the variability of activity patterns across OSNs, is consistent with the possibility that the molecular identity of the odorant receptor expressed by a given neuron is functionally related to the spontaneous activity that neuron displays.

### Uncorrelated spontaneous activity persists in intermediate pupae

Recordings from the developing mammalian (56) and *Drosophila* (49) visual systems have shown that spatiotemporal features of spontaneous neuronal activity change as an animal matures (57, 58). To determine if a similar phenomenon occurs in the developing *O. biroi* olfactory system, we repeated our *in vivo* imaging experiments under identical conditions but with ‘intermediate’ pupae approximately mid-way through metamorphosis (P6 and P7, **Fig. 1 A**). Dramatically more OSNs expressed GCaMP6s in intermediate compared to early pupae **(Fig. 3 A)**, although still visibly fewer than the total Orco-positive population **(Fig. 3 B)**. We manually segmented OSN somata and calculated a ΔF/F timeseries for each cell, resulting in a dataset of 2,225 neurons (445 ± 96.1 neurons per pupa, n = 5 animals; **Fig. 3 C and D**). Like early pupae, OSNs from intermediate animals displayed widespread spontaneous activity, with a ΔF/F event of at least 1 (*i.e.,* a 100% increase) observed in 92.7% of recorded neurons (**Fig. 3 E**). Cells not reaching this threshold were excluded from further analyses. To visualize the distribution of activity types, we again estimated the number of ΔF/F peaks and plotted this estimate against the mean normalized ΔF/F for each cell (**Fig. 3 F**). These distributions overlapped across our two age groups, and we observed a similar variety of activity types in intermediate pupae as we had observed in early pupae (**Fig. 3 G).**

**Figure 3.**
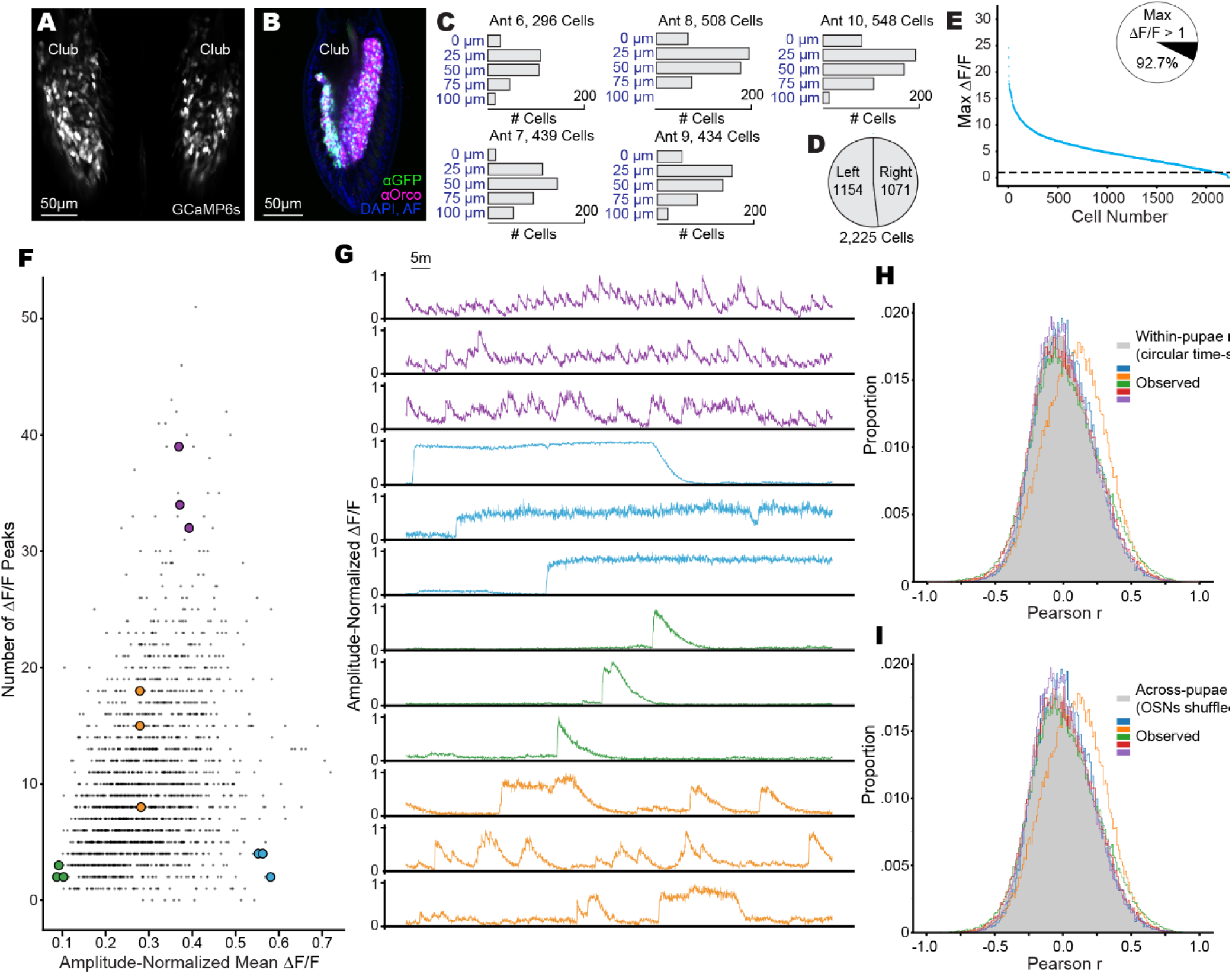
Uncorrelated spontaneous activity persists in intermediate pupae. (A) Two-photon image from a single imaging plane during a volumetric recording session of an intermediate pupa. (B) Confocal micrograph of the antenna of an intermediate pupa. Compared to early pupae, intermediate pupae have more GCaMP-positive OSNs (green), though still fewer than the entire Orco-positive population (magenta). Nuclei are labeled with DAPI, and cuticular autofluorescence (AF) is visible in the same channel (blue). (C) Counts of GCaMP-positive and manually segmented OSN somas recorded using volumetric two-photon microscopy across five imaging planes and five ants. (D) Pie chart showing the total number of OSNs recorded from the left and right antennae across these five individuals. (E) Maximum ΔF/F fold change attained by each neuron in our dataset of 2,225 OSNs. Each point corresponds to a single neuron, and cells are sorted from largest to smallest maximum ΔF/F. The inset shows the percentage of neurons that exhibited at least one GCaMP transient exceeding 1 ΔF/F (*i.e.,* a 100% increase over baseline). (F) Scatterplot comparing the number of ΔF/F peaks with the amplitude-normalized mean ΔF/F. Each data point is one neuron. (G) Representative time-series traces illustrating a diversity of different ΔF/F activity types. The colors of the traces correspond to the colored dots in (F). (H-I) Pairwise correlations between simultaneously recorded OSNs. Histograms compare the observed distribution of Pearson correlation coefficients between OSN pairs to a null distribution generated by circularly shifting the traces in time (H) or shuffling the cells across pupae (I). Each colored line corresponds to the observed data from a different pupa.

We next evaluated the temporal correlation of spontaneous activity in intermediate pupae, using the same approach as for early pupae (**Fig. 3 H and I**). Here, 4 of the 5 pupae displayed little evidence of temporally structured activity across the OSN population, while the fifth animal (“Ant #7”) exhibited a prominent rightward shift – a result that would be expected if GCaMP activity were positively correlated across neurons (orange trace, **Fig. 3 H and I**). Plotting the pairwise Pearson r over time indicated that this correlation was not present throughout the entire trial, but was instead restricted to a large, discrete event during the first 20 minutes of the 2-hour recording (**Fig. S2 A**). Plotting the proportion of neurons engaged in an active episode for each pupa (see methods) showed that Ant #7 began the recording with few active cells but rapidly ramped up as the trial progressed (**Fig. S2 B**). A visual inspection of individual traces from Ant #7 revealed that many cells indeed exhibited a prominent increase in normalized ΔF/F during this ramp (**Fig. S2 C**). This pattern was not observed in any of the other individuals (**Fig. S2 B**).

### Features of GCaMP activity differ between early and intermediate pupae

Aside from this single correlation epoch, dramatic differences in the temporal kinetics of GCaMP6s activity were not immediately apparent between early and intermediate pupae. Still, we reasoned that the calcium signal could vary more subtly across multiple features simultaneously. To test this, we calculated three amplitude-normalized activity metrics for each neuron in both age groups (episode duration, number of peaks, and interpeak interval), sorted the neurons by pupa, and computed a pupa median for each feature. We additionally included a metric that does not depend on our automated estimation of activity episodes and peak locations, the autocorrelation decay constant (τ_auto_), calculated as the lag time at which a cell’s autocorrelation drops below 1/*e*. This analysis revealed that, compared to early pupae, intermediate pupae displayed longer duration activity episodes, fewer peaks, a longer inter-peak interval, and an increased τ_auto_ (**Fig. 4 A and B**). Together, these results indicate that spontaneous OSN calcium kinetics change as pupae mature, with intermediate pupae exhibiting activity that fluctuates on a longer timescale than in early pupae. Differences in the total fraction of time spent in an active episode were modest and marginally non-significant (0.47–0.57 in early pupae, 0.55–0.62 in intermediate pupae, p = 0.056), indicating that the overall level of activity may not differ dramatically with age (**Fig. 4 C**).

**Figure 4.**
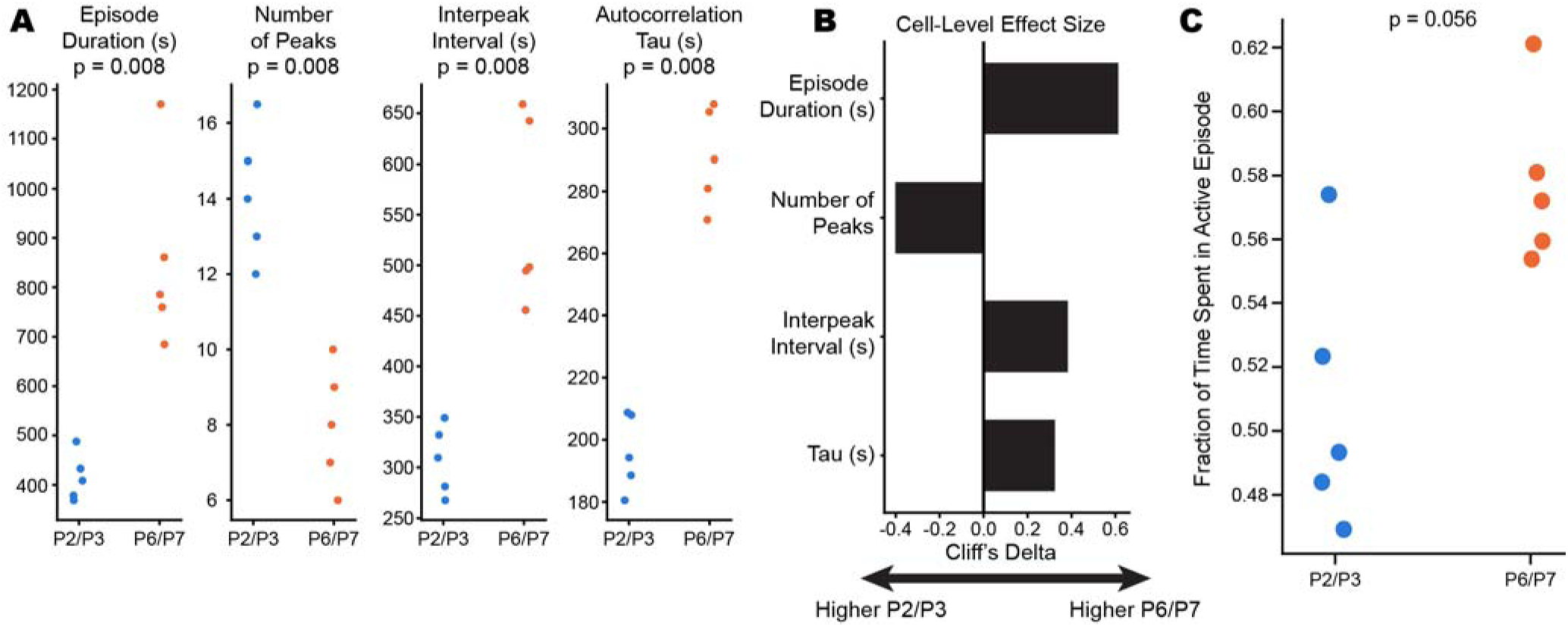
Features of GCaMP activity differ between early and intermediate pupae. (A) Comparison of medians for four activity features across pupal age groups. The measured features are the duration of activity episodes, the number of GCaMP peaks, the interpeak interval, and the autocorrelation decay constant (tau). Statistical significance was determined using Mann-Whitney U tests with p-values indicated above each plot. Note that 0.008 is the smallest attainable p-value. Each data point represents one pupa. (B) Quantification of the cell-level effect size using Cliff’s delta. (C) Median per-pupa fraction of the recording trial spent in an active episode across age groups (Mann-Whitney U test).

### Late pupae show synchronized global calcium events

While only observed in a single trial, the correlated activity bout in Ant #7 suggested that spontaneous activity may become more structured as developing pupae mature. To test this, we extended our recording experiments to a cohort of melanized ‘late’ P16 and P17 pupae, 1-2 days before eclosion (**Fig. 1 A**). Late pupae displayed a global mode of activity not previously observed in any younger ants, where seemingly all OSNs in both antennae rhythmically increased and decreased in GCaMP6s fluorescence in a tightly correlated manner (**Fig. 5A-C**). However, because the high density of GCaMP-positive OSNs in late pupae made it impossible to segment individual somas, we were unable to quantify the precise proportion of neurons participating in these events. To confirm that these global events were not artifacts of infrared laser exposure or mounting the pupa for two-photon microscopy, we observed late pupae under an epifluorescence microscope. Even with this simpler imaging preparation, global calcium events spanning both antennae were clearly visible.

**Figure 5.**
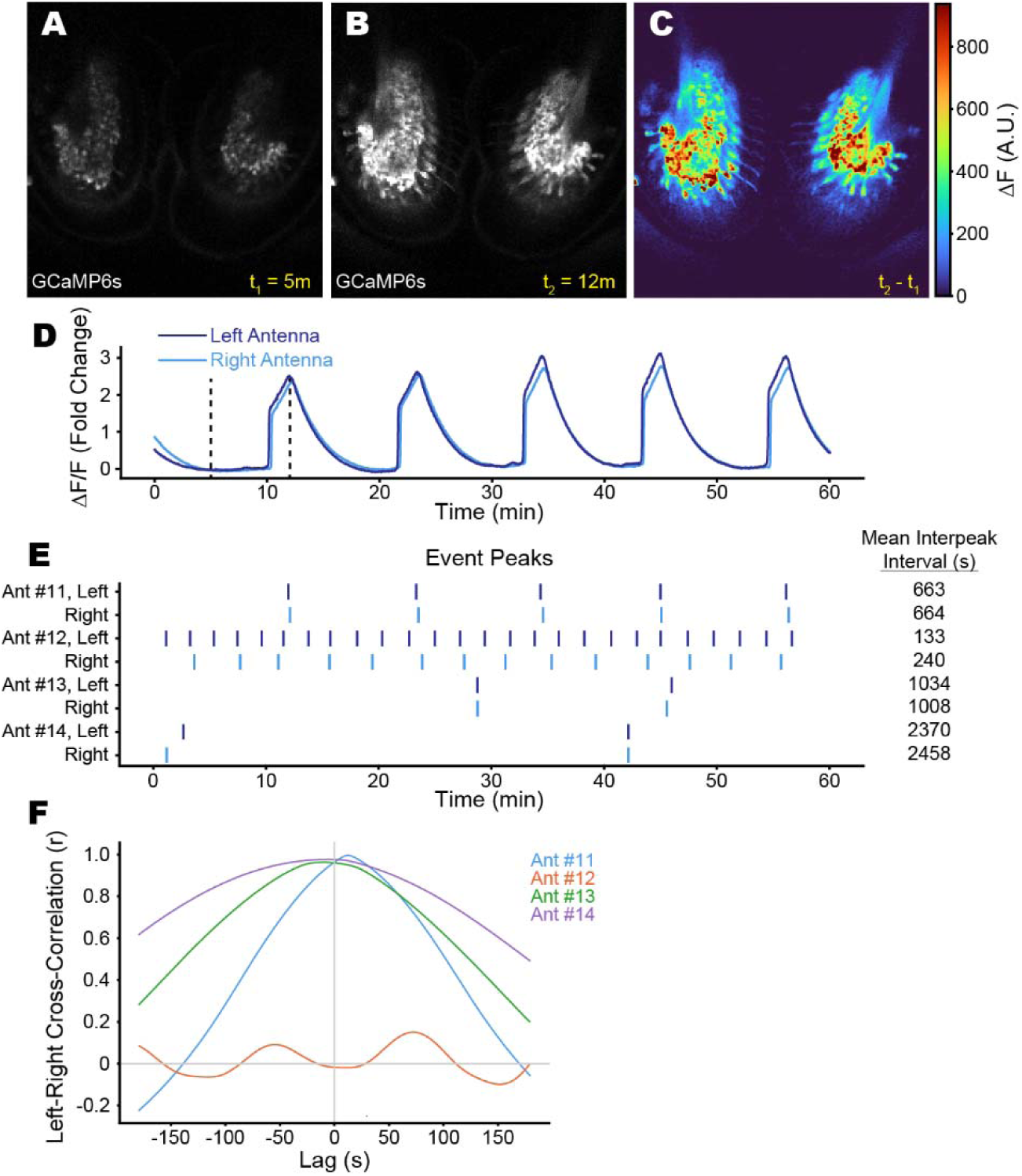
Late pupae show synchronized global calcium events. (A-C) Two-photon micrographs from a recording trial of a late pupa (Ant #11) showing both antennae synchronously increasing in brightness. The frame in (A) was taken 5 minutes after the start of the recording trial (t_1_), and the frame in (B) is from the 12-minute mark (t_2_). (C) Quantification of brightness change displayed as ΔF (t_2_ – t_1_). (D) ΔF/F traces from the entire 1-hour recording trial of the same pupa shown in (A-C). The traces for each antenna are plotted separately in different colors. Vertical, hashed lines correspond to the 5-minute and 12-minute timepoints shown in (A) and (B), respectively. (E) Raster plot showing global event peaks across the dataset (n = 4 pupae). Left and right antennae are plotted in different colors. The mean interpeak interval for each row is indicated on the right. (F) Plot of the cross-correlation coefficient (r) between the left and right antennae at varying lag times, from −180s to +180s, in 1-second steps.

Across animals, global events consistently displayed a sawtooth-like pattern with an abrupt onset across the entire antenna and a slower decay (**Fig. 5D, Fig. S3 A**). However, the period of oscillation of the events differed across ants and, in one case, across the two antennae of the same pupa (**Fig. 5 E and F, Fig. S3 A**). Because of this, the within-antenna mean interpeak interval ranged widely, from 133 seconds to 2,458 seconds – an ∼18-fold difference (**Fig. 5 E**). To quantify the synchrony of the left and right antenna from each ant, we performed a cross-correlation analysis using varying lag times, shifting each ant’s right antenna ΔF/F trace relative to its left antenna trace in one-second steps (±180 s) and computing the Pearson correlation coefficient at each lag. Three of the four animals displayed tight synchrony across the two antennae, indicated by r values peaking near 0s lag time (**Fig. 5 F**). In Ant #12, the two antennae oscillated independently at different periods (133 vs. 240 seconds), indicating that periodicity may be governed independently for each antenna and that bilateral coordination could be a separable process.

## Discussion

In this study, we used *in vivo* two-photon calcium imaging in *Ooceraea biroi* pupae to record the activity of olfactory sensory neurons during metamorphosis. Our experiments revealed that OSN somas are spontaneously active in the absence of olfactory stimuli and that the onset of this activity coincides with the earliest stages of olfactory system assembly (27). Further, we found that the temporal properties of spontaneous activity change as the pupae mature. While early pupae displayed activity with faster kinetics that was uncorrelated across the sensory neuron population, late pupae showed slower activity that culminated in synchronized global calcium events prior to eclosion.

Our results present a striking parallel to spontaneous activity in developing mammalian sensory systems (57). Stimulus-independent activity has been reported across sensory modalities in mammals, and the activity we report here most closely resembles that of the mouse olfactory system. The uncorrelated OSN activity in early pupae mirrors the uncorrelated activity in the neonatal mouse olfactory epithelium (17). In mice, the temporal properties of this activity are determined by which OR a given OSN expresses, and these properties are thought to shape the expression of axon-sorting molecules responsible for normal glomerulus assembly (17). In contrast, spontaneous activity in the developing mammalian visual system has elaborate spatiotemporal structure (59). In the retina, clusters of neighboring ganglion cells fire in bouts that propagate across the retinal surface as waves. These waves advance through developmental stages with distinct properties shaped by different neurotransmitter systems and are required to establish retinotopy in downstream visual areas (57, 60–62). The highly correlated global calcium events we found are reminiscent of these retinal waves, but with the important distinction that they were isolated to late developmental timepoints after all glomeruli had formed and that we did not observe obvious wave-like propagation.

Unlike in mammals, spontaneous activity was long considered dispensable for the development of invertebrate nervous systems. That view has been revised by work in the *Drosophila* pupal optic lobe, where neurons throughout the visual system display patterned, stimulus-independent calcium activity during the second half of metamorphosis that is required for normal circuit formation (49, 58, 63). To our knowledge, however, comparable activity has not been reported in the developing olfactory system of any insect, and the *Drosophila* antennal lobe is thought to assemble without functional ORs or neural activity (19, 27). The activity we describe in ant OSNs also differs from that of the fly optic lobe in two respects. First, it begins far earlier, within the first days of metamorphosis as OSN axons arrive at the antennal lobe, rather than midway through pupal development. Second, it is desynchronized for most of metamorphosis, and only the global events of late pupae resemble the coordinated activity in the fly visual system. The two systems thus appear to recapitulate different mammalian templates, with patterned activity in the fly optic lobe paralleling the retinal waves of the mammalian visual system and the uncorrelated activity of ant OSNs mirroring that of the mammalian olfactory epithelium.

It remains to be determined whether the spontaneous activity we observed is important for the formation of the olfactory map in ants and, if so, whether activity features are instructive for axon targeting or merely permit the survival of nascent olfactory circuits (64). In mice, it is possible to address this question directly using tools that alter OSN excitability (17, 18). Early work established that spontaneous activity is necessary for normal map formation. OSNs expressing the potassium channel *Kir2.1* are delayed in reaching the olfactory bulb and fail to converge on their appropriate glomerulus, instead innervating multiple, disparate glomeruli (18). More recent work argues that spontaneous activity can also be instructive. Optogenetically imposing a different activity pattern on a subset of OSNs expressing the same OR causes stimulated axons to segregate and form additional glomeruli (17). Spontaneous activity in mice thus acts at two levels, permissively by allowing axon outgrowth and targeting to proceed, and instructively by sorting like axons into discrete glomeruli through an OR-dependent temporal activity code. Such a code is an appealing possibility for ants, which, like mice, possess large and rapidly evolving OR repertoires. Each newly duplicated OR must acquire a glomerulus of its own, and a sorting mechanism in which the receptor specifies its own sorting cues would allow new circuits and glomeruli to self-assemble.

Functional experiments similar to those in mice would be instrumental in determining if and how spontaneous activity regulates olfactory development in *O. biroi*, but genetic manipulation of neural activity has not yet been achieved in ants. The clonal raider ant is well poised to change this. Its unique physiology permitted imaging experiments that would not have been possible in other species, and the transgenic protocols that allowed us to produce the *Orco>GCaMP6s* transgenic line could in principle be used to express functional constructs such as *Kir2.1* or optogenetic channels in ant OSNs. The prospect of both imaging and manipulating the olfactory system across metamorphosis positions the clonal raider ant as a genetically tractable model to further investigate how chemosensory circuits develop in an insect with a highly complex olfactory system.

## Materials and Methods

### Clonal raider ant strains and husbandry

Colonies of *Ooceraea biroi* were kept at 25°C in nests consisting of 5cm Petri dishes with a plaster of Paris floor. These nests were watered and colonies were fed frozen fire ant pupae ∼3 times per week during the foraging phase. GCaMP6s ants, which are derived from clonal line B, were propagated by cross-fostering eggs in colonies of clonal line A adults. When GCaMP6s larvae entered metamorphosis, the pupae were transferred alongside an appropriate number of adult workers to a new nest labeled with the date. Three ages of pupae were used in this study: 2-3 day old early pupae, 6-7 day old intermediate pupae, and 16-17 day old late pupae.

### Tissue processing and immunohistochemistry

All ants were anesthetized on ice before dissecting. Antennae from *O. biroi* pupae were removed using microdissection scissors and fine forceps and fixed for 2 hours in 4% PFA in 1X PBS. Antennae were either cut at the scape and processed individually or kept attached to a small piece of head capsule and processed as a pair. After fixation, antennae were blocked for at least 1 hour in 1X PBS containing 5% normal donkey serum and 0.05% Triton-X 100. Samples were incubated in 1:1000 chicken anti-GFP (Abcam, ab13970) and 1:100 mouse anti-Orco (a gift from Vanessa Ruta (12)) primary antibodies for 3 days at room temperature. After washing three times in 1X PBS, the samples were incubated for 2 days in 1:250 donkey anti-chicken Alexa 488 (ThermoFisher, A78948), 1:250 donkey anti-mouse Alexa 647 (ThermoFisher, A32787), and stained with 1:500 DAPI (ThermoFisher, D1306). Antennae were mounted on glass slides in Slowfade Glass mounting medium (ThermoFisher, S36917).

### Fluorescent microscopy

Confocal imaging was performed using Zen image acquisition software on a Zeiss LSM 900 microscope equipped with 405nm, 488nm, 561nm, and 633nm laser lines. A Zeiss LD LCI Plan-Apochromat 40X / 1.2NA objective lens was used with Zeiss Immersol-G immersion medium. For larger samples, multiple image tiles were acquired and stitched in Zen Blue image processing software. Two-photon fluorescence microscopy was performed using a Bruker Investigator microscope equipped with dual GaAsP detectors, a resonant scanning galvanometer, a Z-piezo module for high-speed Z-positioning, PrairieView software, and an Olympus 40X 0.9NA water-immersion objective. Excitation light was provided by a Coherent Axon laser tuned to 920nm and equipped with onboard power control and dispersion precompensation.

### Specimen preparation and in vivo two-photon recording

Prior to recording experiments, a colony containing *O. biroi* pupae was cold anesthetized and adult workers were removed. Anesthetization was only required briefly to separate adults from pupae. To mount pupae for imaging, a small drop of Kwik-Sil surgical silicone adhesive (World Precision Instruments, KWIK-SIL) was used to adhere the dorsal surface of the antennae to a 25mm square glass coverslip. Care was taken to use only the smallest amount of adhesive required to maintain a thin layer between the antennae and coverslip. The adhesive was allowed to cure for 5 minutes, after which the mounted pupa was placed in a humidified Petri dish and allowed to recover for at least 1 hour prior to imaging. For imaging, the coverslip and pupa were removed from the humidified dish and placed in a custom laser-cut acrylic recording chamber with a hole appropriately sized to accommodate the pupa. A piece of damp Kimwipe was added to prevent desiccation. Dental wax was placed at each corner of the coverslip to securely attach the preparation to the acrylic chamber and, using a glass transfer pipette, a drop of deionized water was added to the side of the coverslip opposite the pupa to form a water column with the objective lens. The pupa and deionized water did not come into contact.

Antenna volumes were imaged through an Olympus 40X 0.9NA water-immersion objective lens at 1X optical zoom using a resolution of 512x512x5 voxels (XYZ) with 25µm Z-steps. This resulted in a 295µm x 295µm x 100µm volume. Volumes for early and intermediate pupae were acquired at 1.24Hz (∼0.8 seconds per volume). Late pupae were recorded in a single imaging plane at 1Hz and a resolution of 512x512 or 256x256 pixels. An additional 10-minute buffer was added to the beginning of imaging trials because minor mechanical settling of the preparation often coincided with the onset of laser scanning. This initial period was not included in our analyses.

For explant recordings, antennae from early pupae were removed with microdissection scissors and the exterior cuticle of the antenna was delicately dissected with sharp forceps to expose the underlying neuronal tissue. Explants were transferred to a custom-built acrylic microfluidic recording chamber and pinned to a SYLGARD 184 silicone elastomer (Dow-Corning; World Precision Instruments, SYLG184) substrate using fine tungsten wire (California Fine Wire Co.). The preparation was bathed in an artificial hemolymph (AHL) solution containing the following (in mM): 103 NaCl, 3 KCl, 26 NaHCO_3_, 1 NaH_2_PO_4_, 8 trehalose dihydrate, 10 dextrose, 5 TES, 4 MgCl_2_, and 1.5 CaCl_2_. The solution was continuously bubbled with 95% O_2_, 5% CO_2_ gas and washed over the antenna explant at a rate of 250 mL/hour using a gravity perfusion system. For VUAA1 (Sigma-Aldrich, SML1336) experiments, the recording chamber was fastened to the two-photon microscope stage and explants were washed in control AHL containing 0.5% DMSO. As a precaution, explants were allowed a 5-minute resting period to acclimate to any mechanical disturbances associated with the onset of AHL perfusion. A 10-minute stimulus trial began immediately after this resting period. After 90 seconds had elapsed, a valve was manually triggered that perfused the preparation with a 0.5% DMSO, 100µM VUAA1 AHL solution for the remainder of the 10-minute trial. Each preparation was used only once. Explants were imaged using a single z-plane at a resolution of 512x512 pixels at 1Hz.

### Image processing and data analysis

All image processing and data analysis was performed in ImageJ or using custom Matlab and Python scripts. For all imaging trials, non-rigid motion correction (NoRMCorre (65)) was applied to correct for slight drifts in the XY dimension. For early and intermediate pupae, motion-corrected videos were opened in ImageJ and olfactory sensory neuron somas were manually segmented and tabulated using the ROI Manager plugin. Each ROI included a single neuron soma. We omitted groups of two or more somas immediately adjacent to one another if they could not be separated into distinct ROIs. The GCaMP intensity of each cell was taken as the mean value of all pixels within the ROI. We calculated ΔF/F_0_ traces for each ROI using the formula ΔF/F_0_ = (F – F_0_) / F_0_, where the F_0_ baseline was considered the 12^th^ percentile intensity value for the ROI. A cell was considered active if its ΔF/F_0_ exceeded 1 (*i.e.,* a 100% increase over baseline) for at least 3 consecutive frames (∼2.4s) at any point during the recording trial. Cells that did not meet this criterion were considered inactive and were excluded from further analyses. In total, 11 out of 1,245 cells (0.88%) were excluded in early pupae and 162 out of 2,225 cells were excluded in intermediate pupae (7.28%). Cell counts displayed in Figure 1 and Figure 3 reflect the total number of ROIs prior to exclusion. Amplitude-normalized traces were calculated using the formula ΔF/F_norm_ = (ΔF/F_0_ −ΔF/F_min_) / (ΔF/F_max_ −ΔF/F_min_).

For each pupa, we computed the Pearson correlation coefficient (r) between the raw ΔF/F_₀_ traces of every pair of active cells over the full 2h recording (early group: 1,234 cells across 5 pupae, 156,821 pairs; intermediate group: 2,063 cells across 5 pupae, 444,830 pairs). Pairs on the same or adjacent planes whose ROI polygons passed within 10 pixels of each other were excluded to prevent contamination from passing dendrites and potential segmentation artifacts, resulting in 1.34% and 0.91% of pairs being excluded for early and intermediate pupae, respectively. No cells were entirely excluded. The distribution of r values for each pupa was compared visually against two null distributions, each pooled over 200 shuffle rounds. In the within-pupa circular shift, every cell’s trace was independently rotated in time by a lag drawn uniformly from 5 min to T −5 min (T = recording length), with the displaced segment wrapped to the start of the trace; this preserves each cell’s autocorrelation and amplitude distribution but removes temporal alignment between cells. In the across-pupae shuffle, all cells of an age group were pooled and, in each round, randomly partitioned into five pseudo-pupae matching the cell counts of the real pupae and ignoring true animal identity. Histograms are shown as proportions of pairs per bin; bin width = 0.01.

Active episodes and peaks were detected on smoothed ΔF/F_0_ traces. For each cell, the ΔF/F_0_ trace was smoothed using a second-order Butterworth filter with a cutoff frequency of either 0.02Hz or 0.05Hz. Active episodes were defined on a 0.02Hz-filtered trace to capture sustained states, while peaks were defined on a 0.05Hz filtered trace, which better maintained fast transients. Both peaks and active episodes were defined relative to the standard deviation of each ROI’s per-frame imaging noise (σ), estimated from the unsmoothed ΔF/F_₀_ trace using a median-based approach robust to the presence of calcium events. For each ROI, we computed the frame-to-frame difference over the full recording (Diff = Frame_n_ – Frame_n-1_), took the median absolute deviation (MAD) of these values (MAD = median(|Diff – median(Diff)|)), and calculated σ using the formula σ = MAD / 0.6745 / √2.

Active episodes were defined using a hysteresis-based approach where a trace must exceed a higher threshold to start an episode and fall below a lower threshold to end the episode. This prevents fragmented episodes due to fluctuations around a single value. Thresholds are expressed relative to the cell’s baseline (estimated as the 8^th^ percentile of the ΔF/F_0_). An active episode was detected when the ΔF/F_₀_ exceeded this baseline plus the largest of (i) 4σ, (ii) 10% of the cell’s dynamic range, or (iii) 30% of the previous episode’s peak amplitude. This detection threshold determines whether an episode occurred, but not the initial onset of that episode. After an episode was detected, we defined the onset by searching back from the detection timepoint to the timepoint where the trace first exceeded the baseline by 1σ or 10% of the cell’s dynamic range, whichever value was greater. Episode onset was therefore defined as the moment the trace first departed from baseline, rather than the later moment at which it crossed the higher detection threshold. An episode ended when the trace fell back to within 1σ of baseline or to 20% of the episodes own peak height, whichever was greater. Peaks were found using the SciPy find_peaks function. To be considered a peak, a local maximum was required to have a prominence of at least 3σ. Prominence was evaluated within +/- 10 minutes around each candidate location.

For late pupae, only two ROIs were drawn for each animal (one for the left and one for the right antenna), because segmentation of individual soma ROIs was not feasible due to the high density of neurons at this developmental stage. ΔF/F_0_ was calculated for late pupae using the same method as for early and intermediate pupae. The ΔF image in Figure 5C shows the pixel-wise difference in raw fluorescence between an 11-frame (+/-5s) average at the crest of an event (12min) and an 11-frame average at the preceding trough (5min). Because whole-antenna traces average thousands of pixels, events were detected on the raw ΔF/F_₀_ trace as contiguous runs exceeding the trace minimum plus 30% of its dynamic range, with a minimum duration of 10s.

Event peaks were taken as the maximum ΔF/F_0_ during these windows. Peaks of events already in progress at the start of the recording were excluded, and the mean interval between consecutive remaining peaks is reported per antenna. Bilateral coupling was assessed by computing the Pearson correlation between the left- and right-antenna ΔF/F_₀_ traces at integer lags from −180s to +180s. Positive lags indicate that the left antenna leads.

## Acknowledgments

We thank Stephany Valdés-Rodríguez, Alejandra Hurtado-Giraldo, Alek Rahman, and Leonora Olivos-Cisneros for ant husbandry, as well as Yohann Chemtob for computing assistance. This work was supported by the National Institute on Deafness and Other Communication Disorders under award number K99DC021506 to D.D.F. The content is solely the responsibility of the authors and does not necessarily represent the official views of the National Institutes of Health. This work was also supported by the Howard Hughes Medical Institute via the Investigator program (to D.J.C.K.). This is Clonal Raider Ant Project paper #50.

## Data, Materials, and Software Availability

All original code has been deposited to a GitHub repository (66). Original Python scripts used for statistical analyses and plotting will be made available as a Zenodo repository prior to publication. Functional imaging data and confocal stacks will be made publicly available on the DANDI archive and the Brain Image Library, respectively, prior to publication.

## Author Contributions

D.D.F. and D.J.C.K. conceptualized and designed the study; D.D.F. performed all experiments and analyzed the data; D.D.F. prepared the figures and drafted the manuscript with input from D.J.C.K.; the study was supervised by D.J.C.K. All authors discussed the results, edited the manuscript, and approved the final version.

