## Supplementary figures 1-3 for "Spontaneous neural activity in the developing ant olfactory system"

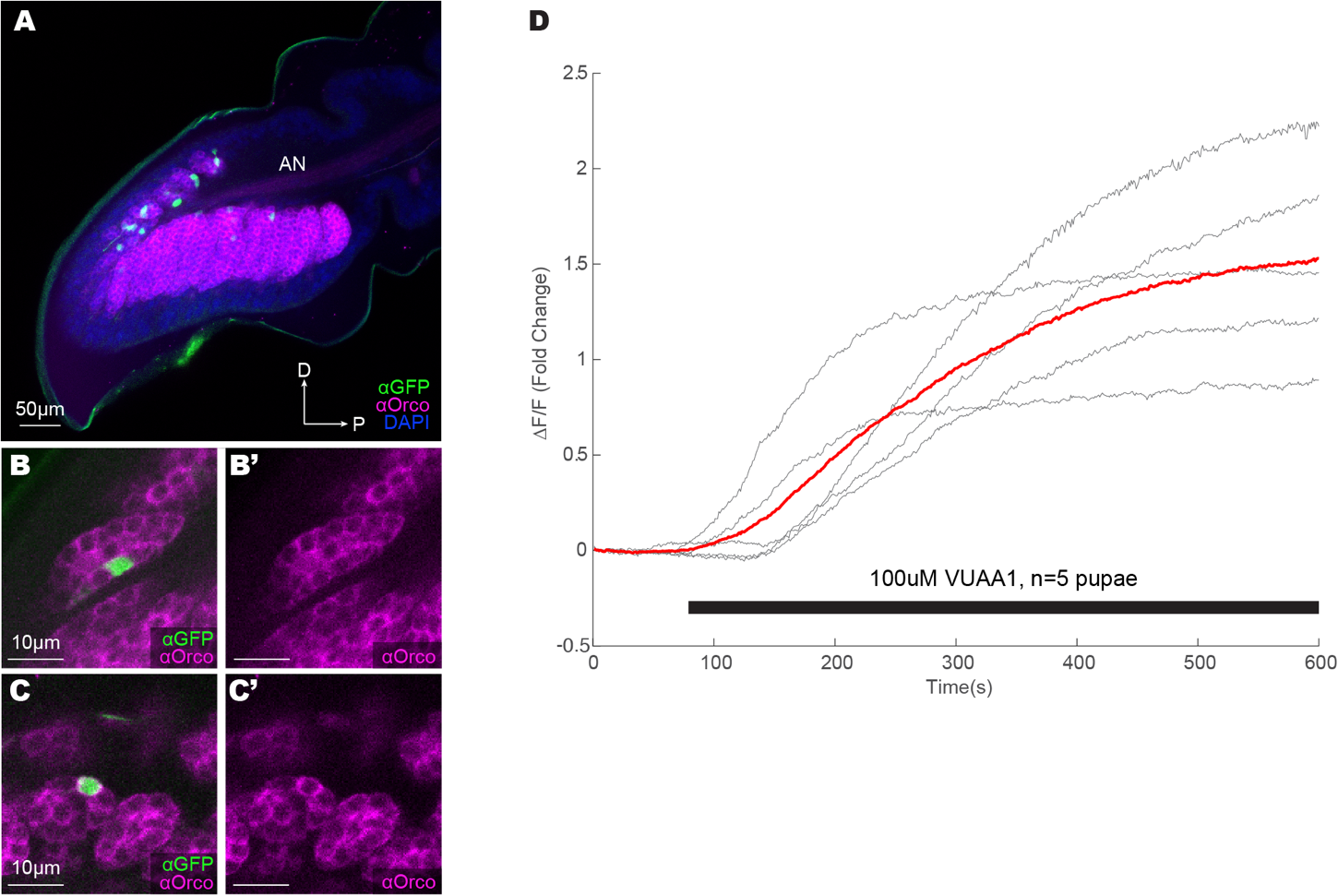

Fig. S1. OSNs in early pupae contain functional odorant receptors. (A) Confocal micrograph of an antenna from a 2-day old pupa, with GCaMP6s-positive (green) and Orco-positive (magenta) cells labeled using immunohistochemistry. Nuclei are labeled with DAPI (blue). (B-C) Two different fields of view comparing the subcellular localization of GCaMP6s and Orco signal. GCaMP6s is cytosolic and fills the soma, while Orco is membrane-bound and can be seen as distinct ‘halos’ at the soma perimeter. (D) Stimulating the antennae of early pupae with VUAA1 in an ex-vivo preparation increases GCaMP6s fluorescence in OSNs. Data were collected from dissected antennae, and 100µM VUAA1 was applied to the bath.

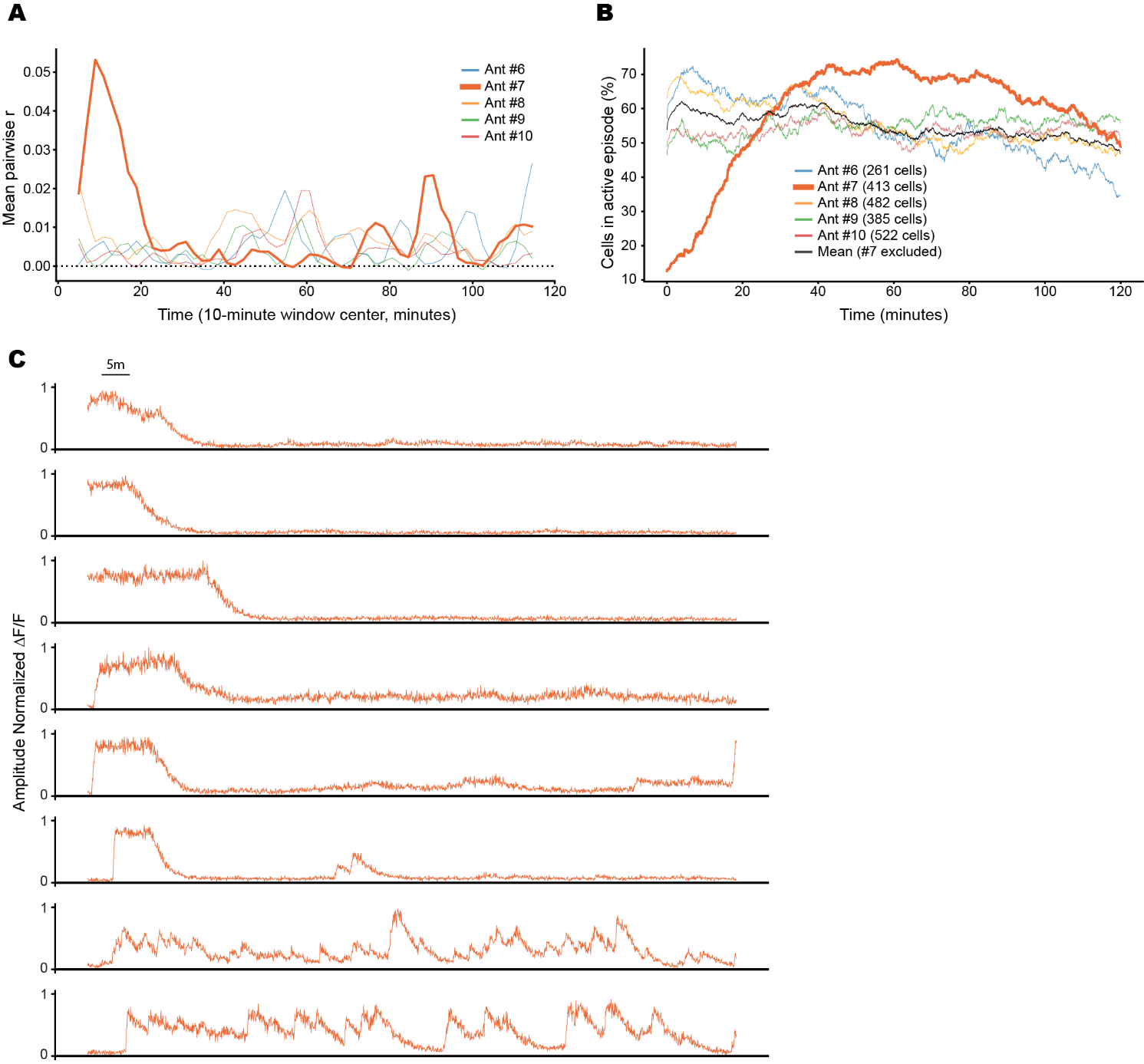

Fig. S2. Analysis of a correlation event in intermediate pupae. (A) Mean pairwise Pearson correlation coefficient (r) plotted across two-hour recording trials. The Pearson correlation coefficient was calculated in 10-minute sliding windows, advanced by 2 minutes. Each colored trace represents a different pupa. (B) Time-series plot showing the percentage of total cells in an active episode. Colored traces are separate pupae, and the black trace is the mean with Ant #7 excluded. (C) A selection of amplitude-normalized ΔF/F traces from Ant #7. Each plot is a different neuron.

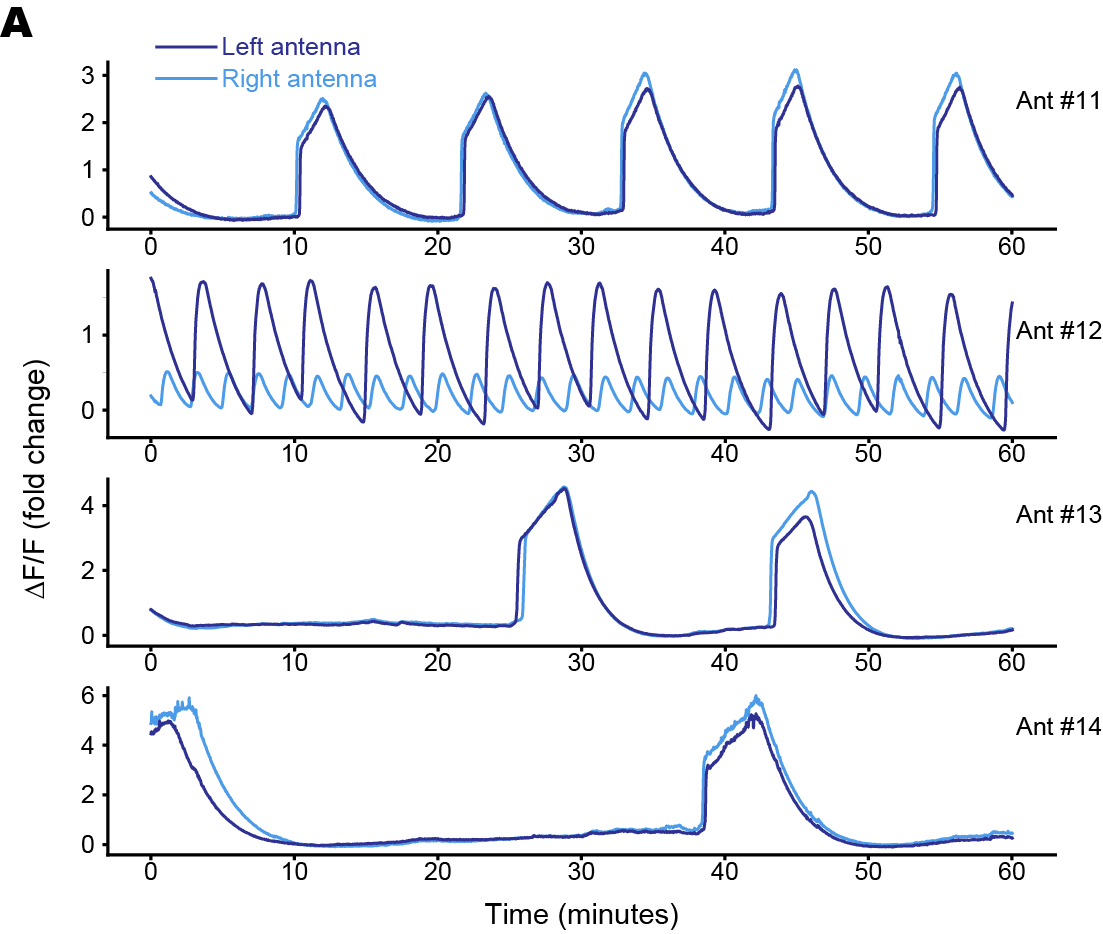

**Fig. S3.** Examples of global calcium events in late pupae. (A) ΔF/F time-series from 4 late pupae during a 1-hour recording session. Data from the left and right antennae of the same animal are plotted as separate traces and labeled in different colors. Each row is a different ant.
